# Stromal netrin-1 coordinates renal arteriogenesis and mural cell differentiation

**DOI:** 10.1101/2023.04.14.536960

**Authors:** Peter M. Luo, Xiaowu Gu, Christopher Chaney, Thomas Carroll, Ondine Cleaver

**Affiliations:** Departments of Molecular Biology, University of Texas Southwestern Medical Center, 5323 Harry Hines Blvd., Dallas, Texas, USA 75390; Internal Medicine and Division of Nephrology, University of Texas Southwestern Medical Center, 5323 Harry Hines Blvd., Dallas, Texas, USA 75390; Center for Regenerative Science and Medicine, University of Texas Southwestern Medical Center, 5323 Harry Hines Blvd., Dallas, Texas, USA 75390

**Author notes:** Present address: Department of Neuroscience, Genentech, South San Francisco, CA 94080. First authors’ and (Peter M. Luo, Xiaowu Gu). Corresponding author email and address: Ondine Cleaver, PhD Department of Molecular Biology, University of Texas Southwestern Medical Center 5323 Harry Hines Blvd., NA8.300, Dallas, Texas 75390-9148, USA.

**Keywords:** Netrin-1, Klf4, artery, mural cell, SMA, NG2, pericyte, nephron, Foxd1

## Abstract

The kidney vasculature has a uniquely complex architecture that is essential to proper renal function. Little is known about the molecular mechanisms that direct where and when blood vessels form during kidney development. We identified a regionally-restricted, stroma-derived signaling molecule, netrin-1 (Ntn1), as a putative regulator of vascular patterning. We generated a stromal progenitor-specific knockout of netrin-1 (*Ntn1^SPKO^*) that resulted in smaller postnatal kidneys with altered epithelial development and profound defects in arterial and capillary architecture. We also found significant loss of arterial vascular smooth muscle cell (vSMC) coverage and ectopic smooth muscle cell deposition at the kidney cortex. Transcriptomic analysis of *Ntn1^SPKO^* kidneys revealed downregulation of Klf4, which we find expressed in stromal progenitors. Deletion of Klf4 in the stroma largely phenocopies loss of Ntn1, and expression of Klf4 in *Ntn1^SPKO^* kidneys rescues ectopic vSMC deposition. Vascular defects observed in *Ntn1^SPKO^* are transient, as both arterial and smooth muscle coverage defects resolve late in development, however ectopic peripheral smooth muscle perdures perinatally. These data suggest a stromal-intrinsic Ntn1-Klf4 axis acting as an essential mediator of stromal crosstalk and vascular progenitor differentiation.

## INTRODUCTION

The vasculature of the kidney is integral to its primary function of removing wastes and extra fluids from the body via urine production. In fact, the kidney filters over 200 liters of blood every single day. This process requires a complicated vascular network, including smaller vessels, such as the glomerular, vasa recta and peritubular capillaries that are designed for solute exchange, as well as a large tree-like arterial network that channels blood throughout the organ (Mohamed and Sequeira-Lopez, 2019). We previously characterized the semi-stereotyped branching of the initial renal arterial tree, which becomes distinguishable as a tributary of the abdominal aorta at 13 days of gestation (embryonic day 13, or E13) in mouse (Daniel et al., 2018). The question arises as to how renal vessels take on the proper architecture that ensures their essential physiological functions. Given that blood vessels are known to respond to paracrine factors from non-vascular cells, it is likely that the various kidney cell types present during development lay down a patchwork of cues that direct vascular patterning. However, to date, these factors remain unknown.

The stroma, or interstitium, of the kidney, plays a crucial role in the development and patterning of kidney structures. Deriving primarily from a population of Foxd1+ progenitors present in the kidney periphery, stromal cells permeate the developing kidney, encasing both epithelial and endothelial structures. Far more than simply providing a supportive matrix for kidney to grow within, the stroma participates in cellular crosstalk that is required for both renal cell fate and overall tissue morphogenesis. (Das et al., 2013; Hurtado et al., 2015; Ide et al., 2018; Mendelsohn et al., 1999; Zhang et al., 2003). Stromal progenitor cells communicate with nephron progenitors (NPCs) to regulate their collective differentiation, and with the ureteric bud (Ub) to direct collecting duct formation and branching (Paroly et al., 2013).

The stroma is also required for proper development of the renal vasculature (Hum et al., 2014), however it is unclear how. Stromal progenitors (SPs) may signal directly to vascular precursors, alternative, they might regulate endothelial cells via their descendants within the perivascular niche. Cortical stromal progenitors give rise to a group of cells collectively termed ‘mural cells’ (Sequeira-Lopez et al., 2015). Mural cells are heterogeneous along the vascular tree, including vascular smooth muscle cells (vSMCs) that tightly and densely enwrap larger caliber vessels, and pericytes that invest more sparsely along capillaries (Armulik et al., 2010; Jain, 2003). Mural cells physically surround and support vascular endothelial cells as they differentiate. In mature vessels, mural cells regulate a host of vascular properties, including vessel patency, quiescence, and tone, as they are exposed to shear stress and pressure from hemodynamic flow. Mural cells have also recently been proposed to regulate the patterning and development of vasculature (Orlich et al., 2022; Stratman et al., 2017).

Development of organ blood vessels is guided by a host of angiogenic factors during development, including the secreted laminin-like protein Netrin-1 (Ntn1). Ntn1 has been shown to be required for the proper development of blood vessels and cell survival (Furtado et al., 2023; Larrivée et al., 2007). Notably, Ntn1 directly impacts endothelial cell migration, function, and survival via signaling to Unc5b (Boyé et al., 2022; Castets et al., 2009; Lu et al., 2004). The role of Ntn1 is not restricted to the endothelial cells, however, as it is also required for axonal migration, lung and pancreatic branching, and other key processes in organ development (Bin et al., 2015; Brunet et al., 2014; Colamarino and Tessier-Lavigne, 1995; Liu et al., 2004; Matilainen et al., 2007; Mediero et al., 2015; Yebra et al., 2003; Yung et al., 2015). Ntn1 mediates these effects via signaling to multiple other receptors, including other members of the Unc5 family, Neo1, and DCC (Freitas et al., 2008; Huyghe et al., 2020).

In this study, we examine the unknown role of Ntn1 in kidney development, with a focus on the renal vasculature. We identified Ntn1 expression in the kidney stroma using single cell data (England et al., 2020) and confirmed this observation *in vivo*. In addition, we show numerous Ntn1 receptors expressed in the kidney, positioning Ntn1 as a potential mediator of stromal crosstalk to other cell types, including the vasculature. Using conditional genetic ablation, we show that Ntn1 is required for proper development of renal arteries and normal distribution of vascular mural cells. Mechanistically, we show that loss of Ntn1 results in the loss of the transcription factor Klf4, which is a key regulator of smooth muscle progenitor fate. Loss of either Ntn1 or Klf4 result in ectopic mural cells in cortical stroma. Together, these novel findings underscore a central role for Ntn1 in kidney development, specifically for patterning and mural cell coverage of the renal vasculature.

## RESULTS

### Foxd1+ stromal progenitors secrete Netrin-1 in the renal cortex during development

To begin investigating the role of netrin-1 (Ntn1) in the developing kidney, we assessed both its transcript and protein expression throughout nephrogenesis. *Ntn1* expression was observed by *in situ* hybridization within stromal progenitors located at the periphery of the cortex from E12.5 to E14.5 **(Fig. S1A-C’)**. Transcript expression was not observed in the medullary region (Med) of the kidney, nor in the nephrogenic zone (NZ), which includes ureteric bud tips and nephron progenitor caps **(Fig. S1C’)**. Ntn1 receptors Unc5c (**Fig. S1D**) and Unc5b (**Fig. S1E**), were expressed in nephron progenitors (NPCs) and blood vessels, respectively, and Neo1 **(Fig. S1F)** was expressed in multiple lineages, including epithelial cells and cortical stroma.

Ntn1 protein, on the other hand, was present within the stroma and amongst NPCs throughout the cortex of the developing kidney **(Fig. 1A-B” and Fig. S1G-J, M, M’)**. Interestingly, we found Ntn1 protein enriched at the tip of the ureteric bud (Ub), where no transcripts are detected via in situ hybridization or single cell analysis **(Fig. S1H**, and data not shown). This observation suggests that the protein found on the UB tips is likely derived from the stromal cells. Over the course of development, Ntn1 protein remained tightly restricted to the nephrogenic zone up to E18.5 **(Fig. S1G-J)**. Protein levels decreased rapidly after birth and were no longer detectable by P3 **(Fig. 1C-C”, Fig. S1K,L**), a timepoint corresponding to the cessation of nephrogenesis. Together, these results support a role for cortical Ntn1 during kidney development.

**Figure 1.**
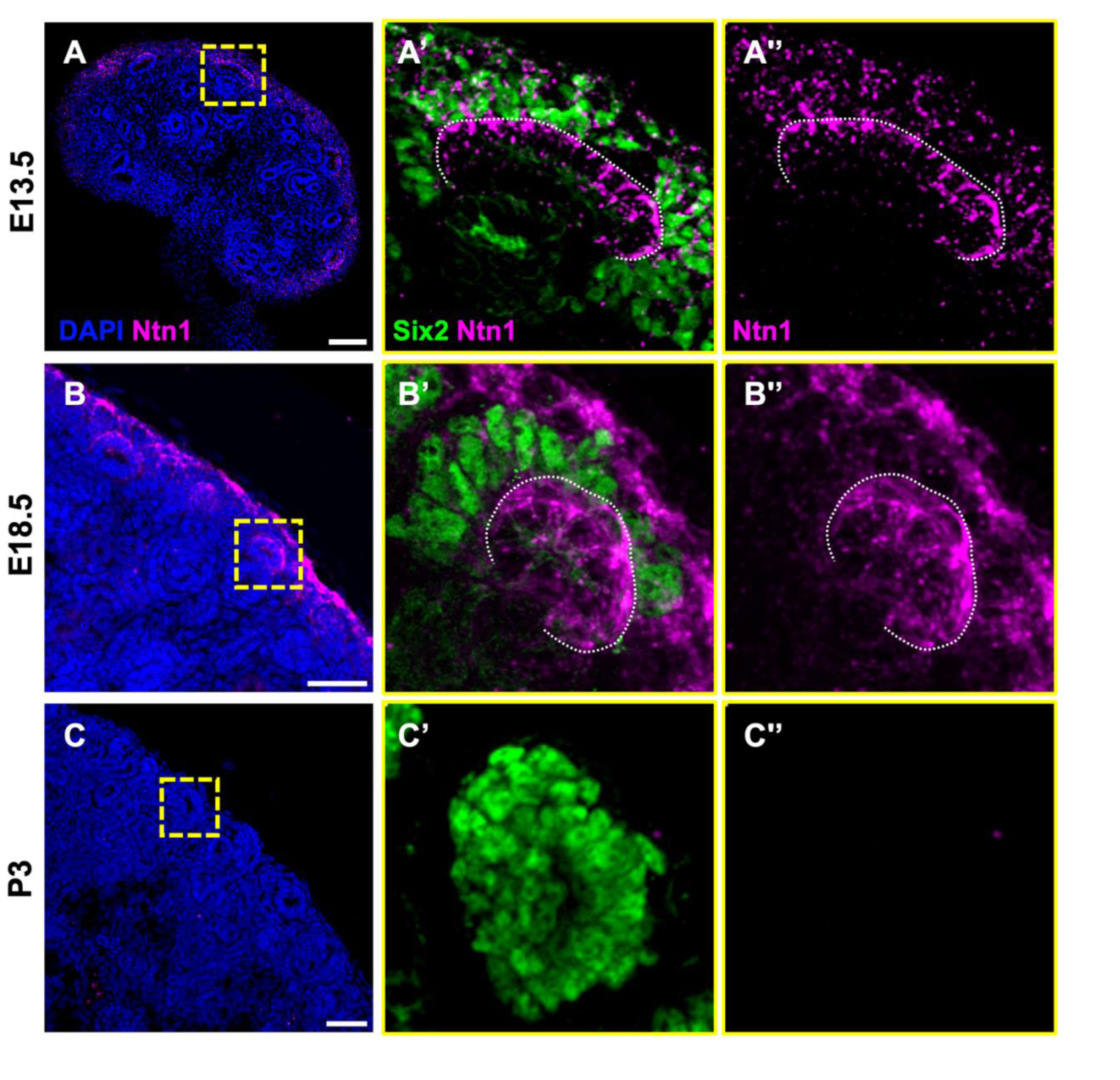
Netrin-1 is expressed by cortical stromal progenitors during embryonic kidney development. (A-C) Immunofluorescence for Ntn1 on kidney sections from E13.5, E18.5, and P3 wild-type embryos, showing cortical enrichment of Ntn1 protein. (A’, A”, B’, B”, C, C”) High magnification insets with costains for Six2 to delineate the nephron progenitor cells, which lie under the Foxd1+ stromal progenitors (not shown) and surround the ureteric bud tip (outlined). Scale bars: 100µm (A), 50µm (B,C).

### Loss of Ntn1 impairs kidney development

Ntn1 is involved in the patterning of the nervous and vascular systems, and its global deletion results in embryonic lethality (Bin et al., 2015; Yung et al., 2015). To test its function in the developing kidney, we generated a kidney stroma specific deletion of Ntn1 (hereafter termed *Ntn1^SPKO^*) by crossing either *Ntn1^f/f^* or *Ntn1^f/+^* females to *Ntn1^f/+^* heterozygote males containing a *Foxd1^GC^* allele **(Fig. 2A)** (Humphreys et al., 2008). We confirmed Ntn1 deletion by both western blot analysis of whole embryonic kidneys **(Fig. 2B)** and immunofluorescence (IF) (**Fig. 2C,D**). Importantly, no Netrin1 protein was detectable in *Ntn1^SPKO^* kidneys, supporting the conclusion that positive staining throughout the nephrogenic zone was derived from stromal progenitors.

**Figure 2.**
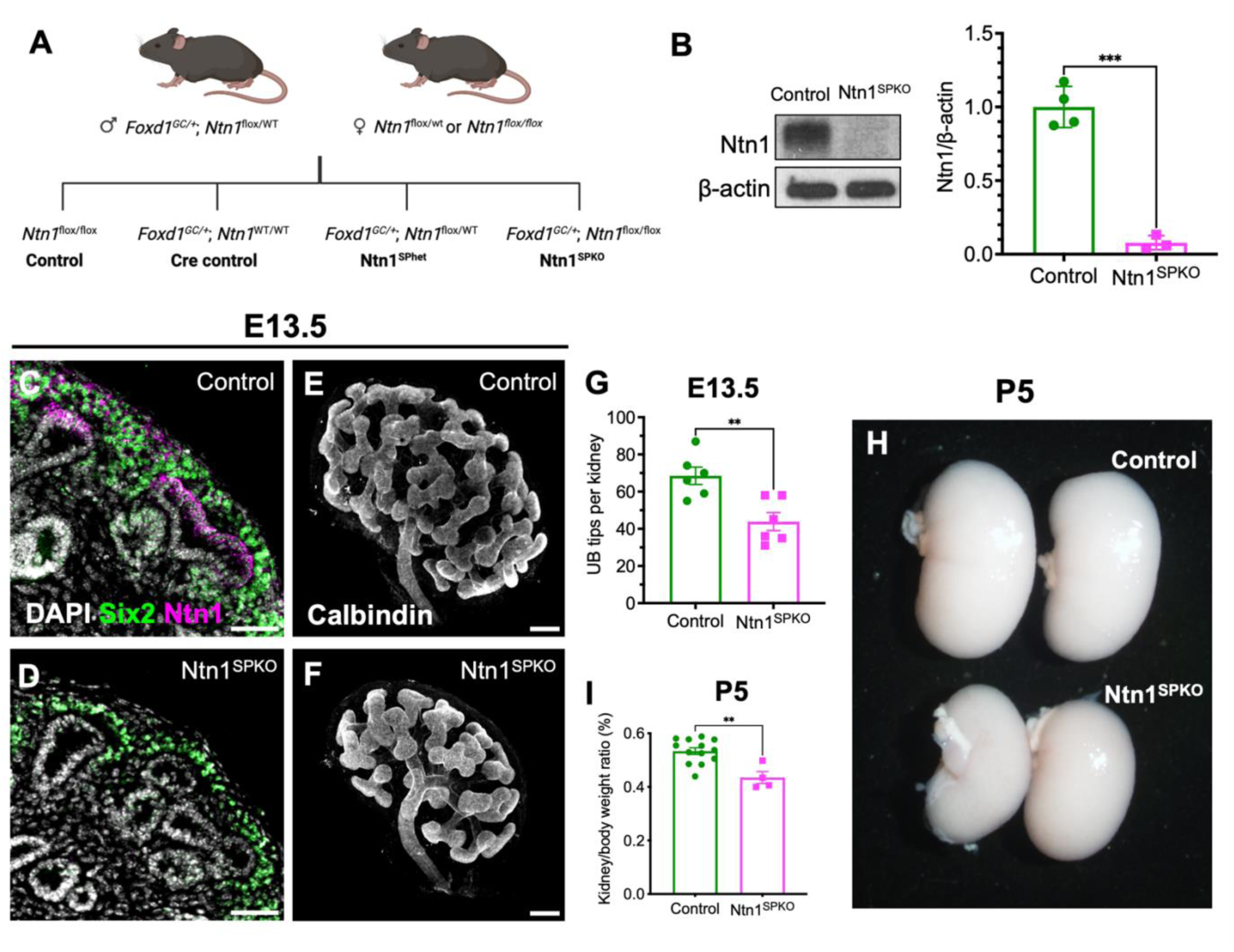
Ablation of netrin-1 in the kidney stroma impairs kidney development. (A) Breeding scheme of stromal progenitor specific knockout of Ntn1 (*Ntn1^SPKO^*) using the Foxd1-Cre. (B) Western blot analysis of Ntn1 levels in *Ntn1^SPKO^* kidneys showing complete loss of protein (n=3, p=0.0001). (C-D) Immunofluorescence for Ntn1 and Six2 on E13.5 kidney sections, showing loss of Ntn1 protein and normal nephron progenitor cell morphology in Ntn1^SPKO^. (E-G) Whole mount immunofluorescence (WMIF) of E13.5 kidney ureteric tree with calbindin, showing smaller kidneys and decreased branching quantified by number of ureteric bud tips (n=6, p=0.0044). (H) Photo of P5 kidneys, showing smaller *Ntn1^SPKO^* kidneys after cessation of nephrogenesis. (I) Kidney weight/body weight comparison demonstrates decreased kidney size in *Ntn1^SPKO^* embryos (n=4, p=0.0016). Scale bars: 50µm (C-D), 50µm (E,F).

Grossly, *Ntn1^SPKO^* kidneys were smaller than those of littermate control embryos during development **(Fig. 2E,F).** To assess specific defects in kidney development, we analyzed Ub branching in E13.5 *Ntn1^SPKO^* kidneys by whole mount IF for Calbindin and observed significantly fewer epithelial tips **(Fig. 2E-G)**. Notably, Six2+ NPCs remained present in the nephrogenic zone without Ntn1 and appeared mildly reduced **(Fig. 2C,D)**. Staining for DBA and Pax8 at P5 showed relatively normal morphology of ureteric and nephrogenic epithelia, respectively **(Fig. S2A-B).** However, Six2+ NPCs were observed in *Ntn1^SPKO^* kidneys as late as P5, a time point at which control kidneys experience cessation of normal nephrogenesis and depletion of NPCs **(Fig. S2C-E)**. Perduring Six2+ cells in *Ntn1^SPKO^* kidneys also continued to express NCAM, a cell adhesion molecule that indicates progression of differentiation towards the renal vesicle stage of nephrogenesis **(Fig. S2C-D’)**. These data suggest that in the absence of Ntn1 a proportion of NPCs differentiates and undergoes MET, yet are maintained past the point when they are normally depleted.

The reduction of kidney size in mutants lasted after development. This was evident by direct comparison of P5 WT and mutant kidneys **(Fig. 2H)**, but also when taking into consideration kidney to body weight ratios **(Fig. 2I)**. To assess the effect of Ntn1 loss on nephron generation, we performed acid maceration of adult kidneys followed by glomerular counting and found significantly fewer glomeruli in *Ntn1^SPKO^* kidneys **(Fig. S2F).** Glomerular counts correlated with kidney/body weight ratio, suggesting that this decrease was in line with the observed decrease in kidney size **(Fig. S2G)**. These data suggest that the prolonged postnatal expression of Six2 does not result in an overall increase in the number of nephrogenesis cycles, but instead represents perdurance of progenitors following failed nephron differentiation.

### Impaired arteriogenesis and vascular patterning occurs in the absence of Ntn1

Ntn1 is known to regulate angiogenesis and to pattern the vasculature of other organs (Larrivee et al., 2007; Park et al., 2004; Wilson et al., 2006). We asked whether Ntn1 might play a role in the development of the renal arterial tree. Immunostaining for mature arteries with connexin 40 (Cx40, or Gja5) showed drastically reduced expression within *Ntn1^SPKO^* kidney vessels **(Fig. 3A,B).** Expression of a broader arterial marker, Nrp1 **(Fig. 3C,D)** (You et al., 2005) revealed that arterial structures were present within the kidney, although they were grossly mispatterned. Given that Cx40 is normally expressed in arteries in response to blood flow (Chong et al., 2011), we posit that mispatterning of arterial structures in *Ntn1^SPKO^* kidneys may lead to decreased blood flow into the kidney and sought to pinpoint how.

**Figure 3.**
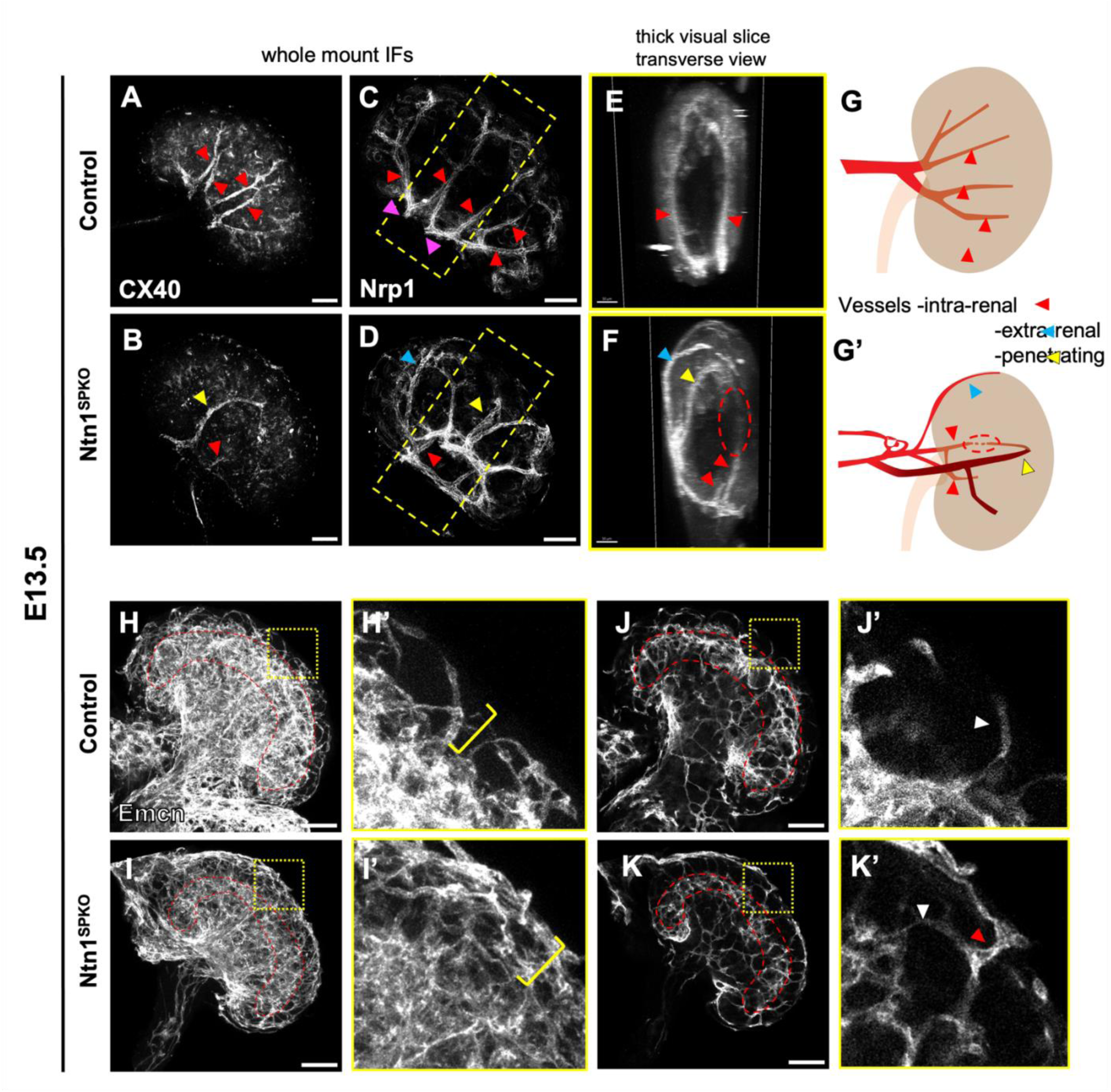
*Ntn1^SPKO^*results in altered patterning of early renal vasculature, including loss of intra-renal arteries, and gain of ectopic arteries and cortical capillaries. (A-B) WMIF of E13.5 kidneys for Cx40, showing fewer mature arteries that have remodeled under flow. Yellow arrowhead indicates a vessel entering the kidney from the cortex rather than the hilum (‘penetrating’), red arrowhead indicates renal artery entering at hilum (‘intra-renal’) with less Cx40. (C-D) WMIF of arterial marker Nrp1 in E13.5 kidneys, showing altered patterning without Ntn1. Yellow and red arrowheads continue to indicate penetrating and intra-renal arteries, respectively. Blue arrowhead indicates artery travelling along ventral kidney surface (‘extra-renal’). (E-F) 200µm slice, viewed orthogonally, through center of Nrp1 WMIF, showing arteries fully within control kidneys and penetrating and extra-renal arteries in *Ntn1^SPKO^* kidneys (yellow, blue arrowheads). Intra-renal arteries (red arrowheads) fail to progress in *Ntn1^SPKO^* kidneys but likely connect to penetrating arteries (dotted circle). (G) Model of arterial patterning defects in *Ntn1^SPKO^* kidneys, showing intra-renal, extra-renal, and penetrating arteries. (H-I) WMIF for endomucin marking capillary vasculature in E13.5 kidneys, showing scant capillaries outside the cortical vascular plexus (red outline) in control kidneys and a thick mesh of capillaries at the surface of the kidney and nephrogenic zone (inset H’-I’, yellow brackets) in *Ntn1^SPKO^*. (J-K) Single slice images of endomucin WMIF showing cortical vascular plexus (red outlines) and vessels along the kidney surface in *Ntn1^SPKO^*. (J’-K’) insets showing small caliber vessels originating from the cortical plexus in both control and *Ntn1^SPKO^* kidneys (white arrowhead), but larger capillaries travelling along surface of *Ntn1^SPKO^* kidneys (red arrowhead). Scale bars: 100µm (A-D, H-K), 50µm (E-F).

Comparing arteries in *Ntn1^SPKO^* and control kidneys, we noted significant differences in vascular patterning. Wild type renal arteries form a ramifying tree-like structure. This tree consists of two main arms (magenta arrowheads) that enter the kidney centrally at the hilum and branch regularly into the upper and lower pole respectively, while remaining within the bounds of the kidney **(Fig. 3C,E,G**, red arrowheads). By contrast, extra-renal arteries in mutants were observed traveling along the surface (blue arrowheads), sometimes looping and entering the kidney directly through the cortex to the middle of the kidney (termed ‘penetrating arteries, yellow arrowheads), as opposed to at the hilum in controls **(Fig. 3D,F,G)**.

To assess where these vessels originated, we performed immunostaining on whole abdominal arterial systems to define the path of the renal artery from the aorta. In control embryos, the renal artery forms a single large branch from the midline dorsal aorta and invades the kidney via the hilum **(Fig. S3A,** grey arrowhead). In *Ntn1^SPKO^* embryos, we observed premature branching and formation of an “arterial plexus” (yellow arrowheads) between the aorta and the kidney, resulting in misdirection of arteries that bypass the hilum and enter the kidney via the cortex (blue arrowheads) **(Fig. S3B)**. We also observed multiple aortic branches contributing to the renal vasculature via collaterals as opposed to only one in control kidneys **(Fig. S3B**, white arrowheads).

Arteries in *Ntn1^SPKO^* kidneys that did enter the kidney at the hilum (‘intra-renal’ arteries) did not progress normally into the kidney **(Fig 3D,F,G, S3D,** red arrowheads). In some cases, penetrating arteries looped and connected with intra-renal arteries **(Fig. 3F,G**, red dotted circle between yellow and red arrowheads). We note that penetrating arteries and their looping morphology are visible by stronger Cx40 staining than intra-renal arteries in *Ntn1^SPKO^* kidneys, suggesting that flow may be shunted away from the hilum at the arterial plexus **(Fig. 3B**, yellow arrowhead). In addition, we observed ectopic collateral connections between intra-renal arteries near the hilum of *Ntn1^SPKO^*kidneys **(Fig. S3D, D’,** orange arrowheads), as opposed to only at the cortex via arcuate arteries in control kidneys **(Fig. S3C, C’,** white arrowheads). These connections often crossed the midline of the kidney, connecting the two main renal artery branches improperly and potentially limiting flow into the kidney.

To assess arterial fate in mutant vessels, WMIF for Sox17 was carried out in control and *Ntn1^SPKO^* kidneys. While expressed at lower levels, arterial endothelial cells in *Ntn1^SPKO^* kidneys expressed detectable Sox17, indicating that arterial identity was not completely disrupted in the absence of Ntn1 **(Fig. S3E,F).**

Next, we analyzed whether arterial mispatterning perdured or progressed as the kidney developed with WMIF for smooth muscle covered-arteries with alpha smooth muscle actin (aSMA). In E15.5 Ntn1^SPKO^ kidneys, we continued to observe penetrating and extra-renal arteries **(Fig. S3G-H,** yellow and blue arrowheads). We notice that penetrating arteries enter the kidney much closer to the hilum compared to E13.5, sometimes making complete turns from their initial direction to direct blood towards the cortex **(Fig. S3H’)**. This is likely due to growth of the kidney, resulting in a shorter relative distance from the point of entry of arteries to the hilum and a gradual restoration of overall corticomedullary patterning. As at E13.5 **(Fig. 3D,F,G**, yellow arrowheads), E15.5 penetrating arteries also connected via a looping structure resembling an arcuate artery to the vasculature within the kidney (**Figure S3H,** red circle). We speculate that Ntn1 does not direct overall corticomedullary direction of arterial growth, but rather general arterial patterning and branching. At this later stage, connecting collateral arteries remained present around the hilum **(Fig. S3H”,** orange arrowheads) and were covered in smooth muscle (aSMA), but closer analysis with Sox17 immunostaining showed abnormal extra branches of these collaterals that were not covered in smooth muscle **(Fig. S3H”-H”’,** red arrowheads).

By E18.5, arterial mispatterning in *Ntn1^SPKO^* kidneys largely resolve. Analysis using staining for aSMA showed a relatively properly branched tree, with only a few abnormal arteries (white arrowheads) remaining at the hilum of the kidney **(Fig. S3I-J’’)**. Together, these findings suggest that extrarenal arteries and other abnormal branches are not covered by smooth muscle and may be pruned due to their potential immaturity.

### Loss of netrin-1 results in aberrant capillary vasculature in the nephrogenic zone

In addition to the arterial tree, which is thought to arise via direct branching of the aorta, the early kidney is populated with a dense vascular plexus that likely arises from endothelial progenitors within the metanephric mesenchyme (Daniel et al, 2018). At E13.5, these fine capillaries form a cortical plexus at the edge of the cortex beneath the nephrogenic zone, wrapping around the entire kidney. The nephrogenic zone itself is relatively avascular during early kidney development (Daniel et al., 2018).

We analyzed the formation of the vascular plexus and capillary organization in *Ntn1^SPKO^* kidneys by performing immunostaining for endomucin **(Fig. 3H-K)**. While in control kidneys, the nephrogenic zone is relatively devoid of vessels **(Fig. 3H,H’**, yellow bracket, **J,J’)**, the cortex of *Ntn1^SPKO^* kidneys is densely packed with capillaries **(Fig. 3I,I’**, yellow bracket). Single visual slices of whole mount immunostaining show that vessels entering into the nephrogenic zone from the cortical vascular plexus in control kidneys are smaller in caliber **(Fig. 3J’**, white arrowhead), whereas larger caliber external vessels are seen at the surface connecting to smaller internal vessels coming directly off of the cortical vascular plexus **(Fig. 3K’**, white and red arrowheads). It is possible that these surface associated capillaries also arise from outside of the kidney and connect to the internal vasculature of the kidney similar to the arteries.

### Loss of renal netrin-1 impairs recruitment of arterial smooth muscle and premature differentiation at the surface of the kidney

It has long been known that recruitment of mural cells, including vSMCs, is an important step of arterial development and maturation (Gaengel et al., 2009). Because mural cells derive from the cortical stromal progenitors (i.e. the Foxd1+ lineage) in the kidney (Humphreys et al., 2010), we aimed to characterize whether stromal Ntn1 ablation mural cell recruitment. We assessed vSMC localization in control and *Ntn1^SPKO^* kidneys by adjacent localization of aSMA and Sox17 immunostaining in E13.5 kidneys. We found that *Ntn1^SPKO^* kidneys displayed significantly fewer arteries covered by vSMCs **(Fig. 4A-C)**. Arteries in control kidneys were covered with aSMA+ cells along their length, well into the kidney, while *Ntn1^SPKO^* kidneys showed little coverage **(Fig. 4A’,B’**, and insets **A”,B”)**. Instead, we found aSMA+ cells crowded the surface of the kidney, where cortical stromal progenitors reside (insets **A’’’,B’’’)**.

**Figure 4.**
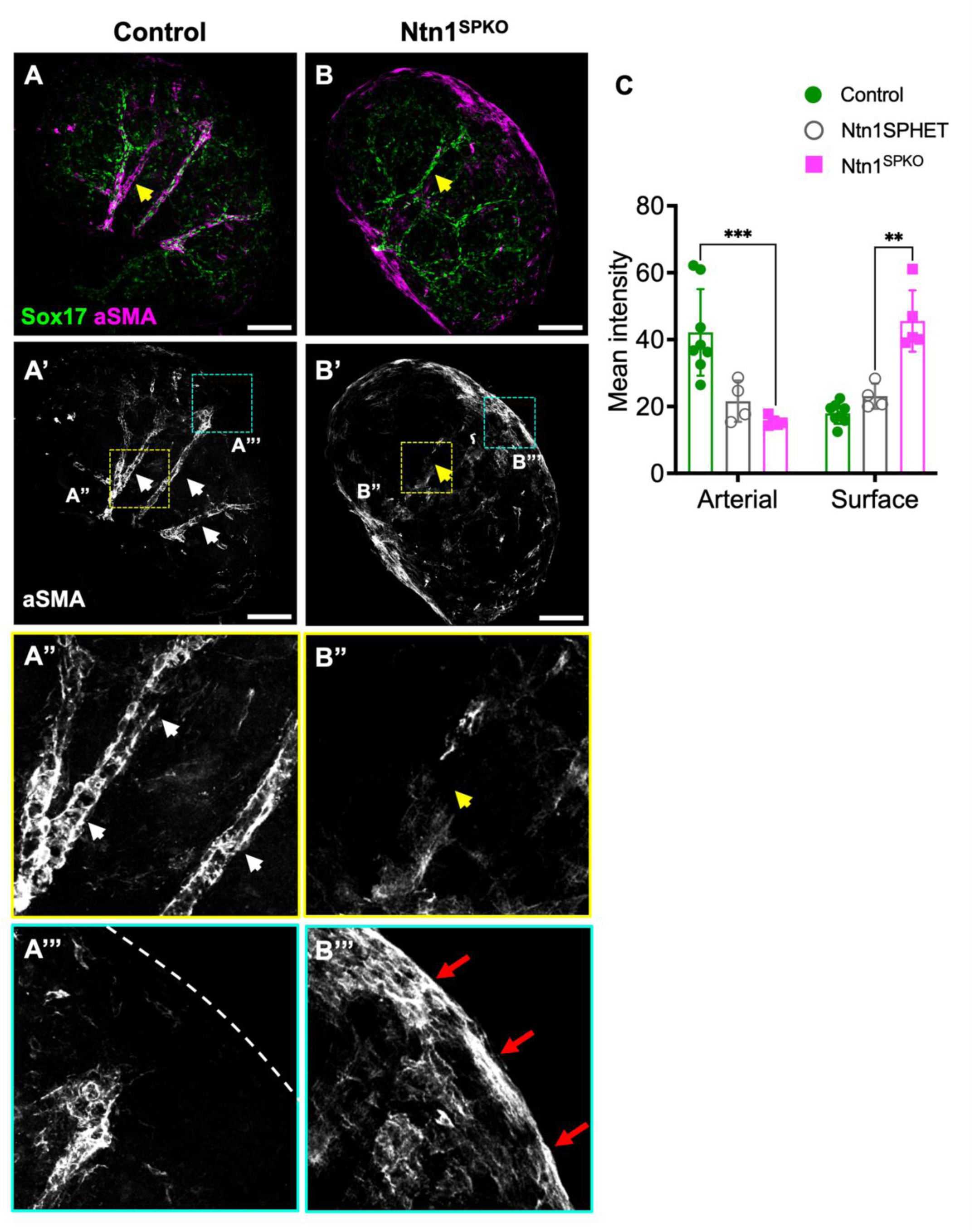
*Ntn1^SPKO^* kidneys exhibit defective vSMC distribution at E13.5. (A-B) WMIF of arterial marker Sox17 and smooth muscle marker aSMA in E13.5 kidneys, showing improper distribution of smooth muscle cells in *Ntn1^SPKO^*. Single channel images (A’-B’) and insets (A’’-B’’) show decreased smooth muscle at arteries (yellow arrowheads) and ectopically differentiated smooth muscle cells at the kidney surface (red arrows). Dotted line in control indicates kidney surface. (C) Quantification of arterial and surface smooth muscle coverage in control vs *Ntn1^SPKO^* kidneys, measured by mean pixel intensity within 25µm or 40µm of the Sox17+ arteries or kidney surface, respectively. (n=5, p=0.000881 (arterial), 0.0027 (surface)). Comparison of *Ntn1^SPHET^* (*Foxd1^GC/+^;Ntn1^f^*^/+^) and *Ntn1^SPKO^* trends towards significance (0.0662). Scale bars: 100 µm (A-B).

To confirm that ectopic SMCs are derived from the same progenitors that contribute to normal vSMCs, we crossed in a fluorescent reporter allele to the *Foxd1^GC^*line to visualize the lineage of Foxd1+ stromal progenitors in the mutants. We found that at E13.5, ectopic smooth muscle cells and normal arterial vSMCs were indeed derived from Foxd1 expressing stromal progenitors **(Fig. S4A-B”)**.

Notably, at E15.5, arteries were covered with smooth muscle **(Fig. S4C, D)**. When examining the coverage distribution of aSMA along arteries, we found that first order vessels (largest arteries entering the kidney from the hilum, green arrowheads) showed equivalent levels of expression, but second order vessels (branches off the first order branches, including arcuate arteries, yellow arrowheads) showed significantly less coverage in *Ntn1^SPKO^* **(Fig. S4C,D**, quantification in **Fig. S4E, and Fig. S4F-G’)**. We noted that when present, penetrating arteries had stronger staining that decreased as they looped and connected with vessels within the kidney. Ectopic aSMA+ cells, however, remained present at the surface of the kidney **(Fig. S4F”,G”** yellow arrowheads), suggesting that eventual coverage of arteries by smooth muscle and ectopic accumulation of smooth muscle at the surface are separate, but related, processes. Together these results underscore the requirement for Ntn1 for normal distribution and investment of vSMCs.

### Netrin-1 is required to suppress NG2+ cell localization at the kidney periphery

Given that stromal-derived mural cells include both vSMCs and pericytes, we asked whether pericyte distribution was affected in the absence of Ntn1. A marker often used for identifying pericytes is NG2, which while not specific to pericytes is useful when combined with cellular localization of positive cells around vessels and capillaries. We therefore interrogated recruitment of pericytes to smaller vessels by performing immunostaining for NG2 and endomucin in E13.5 kidneys. Surprisingly, we found that at E13.5, NG2 staining was not present around capillaries in either control or *Ntn1^SPKO^* kidneys **(Fig. 5A-B’).** Instead, we found that NG2+ cells were present around larger arteries in both control and mutant vessels (**Fig. C-D’**). We noted that costaining with aSMA in control kidneys showed that smooth muscle cells closest to the arteries costained for NG2, but outer layers of mural cells exhibited NG2 expression, with little to no aSMA **(Fig. 5C’**, white arrowhead). NG2 expression around arteries within *Ntn1^SPKO^*kidneys was lower but visible, indicating that some NG2+ mural cells were still able to invest arteries **(Fig. 5D’**, red arrowheads). Notably, penetrating arteries had more NG2 staining, again suggesting that these vessels may have more mural cell investment from early stages **(Fig. 5D’**, yellow arrowhead). However, NG2+ cells were also visible at the surface of the kidney **(Fig. 5E**, green arrow). Interestingly, costaining with aSMA (magenta arrow) showed a lack of colocalization with ectopic smooth muscle at the surface, suggesting that the two cell types are distinct.

**Figure 5.**
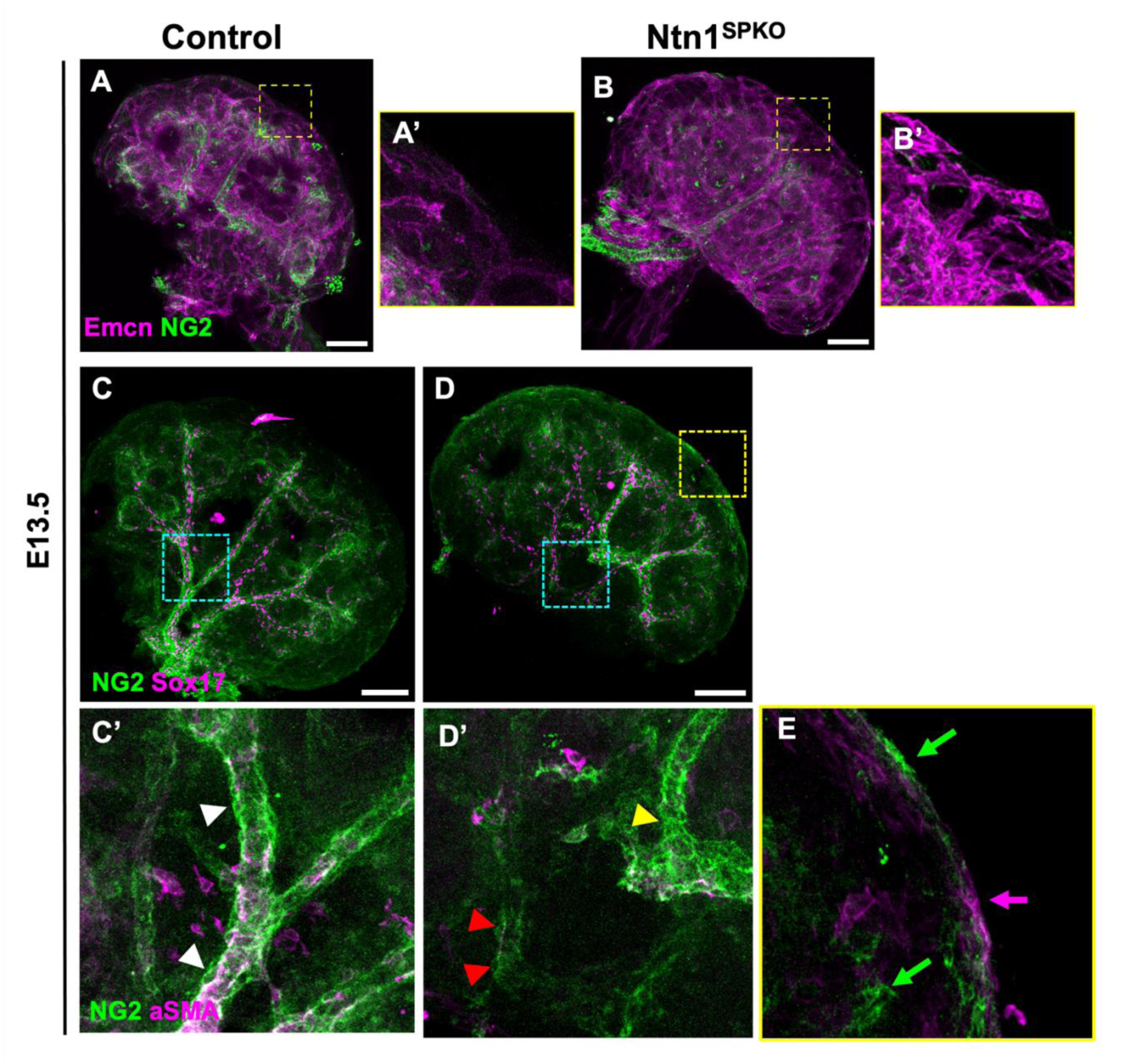
Loss of Ntn1 from the kidney stroma results in ectopic NG2+ cells at the kidney surface and decreased investment along arteries. (A-B) WMIF for NG2 and endomucin, showing no pericyte investment to capillaries beyond the cortical vascular plexus (inset, A’,B’) by E13.5 in either control or *Ntn1^SPKO^*kidneys. (C-D) WMIF for NG2 and Sox17, showing NG2 expression around arteries in E13.5 control kidneys and altered localization in *Ntn1^SPKO^* kidneys. Inset around intrarenal arteries (C’-D’) with costains for aSMA show NG2+/aSMA-mural cells around arteries in control kidneys (white arrowheads), and decreased NG2 coverage of intra-renal arteries in *Ntn1^SPKO^*. Yellow arrowhead indicates penetrating artery with strong NG2 staining. Inset at surface of *Ntn1^SPKO^* kidney (E) shows both NG2+ cells (green arrow) and aSMA+ cells (magenta arrow) at surface, but lack of overlap in expression. Scale bars: 100µm (A-D).

Later in development, at E18.5, NG2+ pericytes were found to associate normally with capillaries in both control and *Ntn1^SPKO^* kidneys, both in the kidney cortex in glomeruli and peritubular capillaries **(Fig. S5A,B),** and in the medulla **(Fig. S5C,D)** around descending vasa recta.

### Loss of netrin-1 results in transcriptional dysregulation of vascular smooth muscle phenotypic switch machinery via loss of Klf4

To determine the mechanisms by which loss of Ntn1 affects developing kidneys, we sought to identify transcriptional changes in *Ntn1^SPKO^*kidneys. We isolated RNA from E13.5 *Foxd1^GC^* control and *Ntn1^SPKO^*mutant kidneys and performed whole transcriptome bulk RNA sequencing, followed by differential gene expression analysis. When compared to controls, 193 genes were found to be significantly downregulated (p<0.1), and 26 genes were significantly upregulated in *Ntn1^SPKO^* kidneys **(Fig. 6A)**. Gene set enrichment analysis showed downregulation of developmental pathways in the absence of Ntn1, especially those involved in muscle cell differentiation **(Fig. 6B)**. The transcription factor Klf4 was amount these downregulated genes.

**Figure 6.**
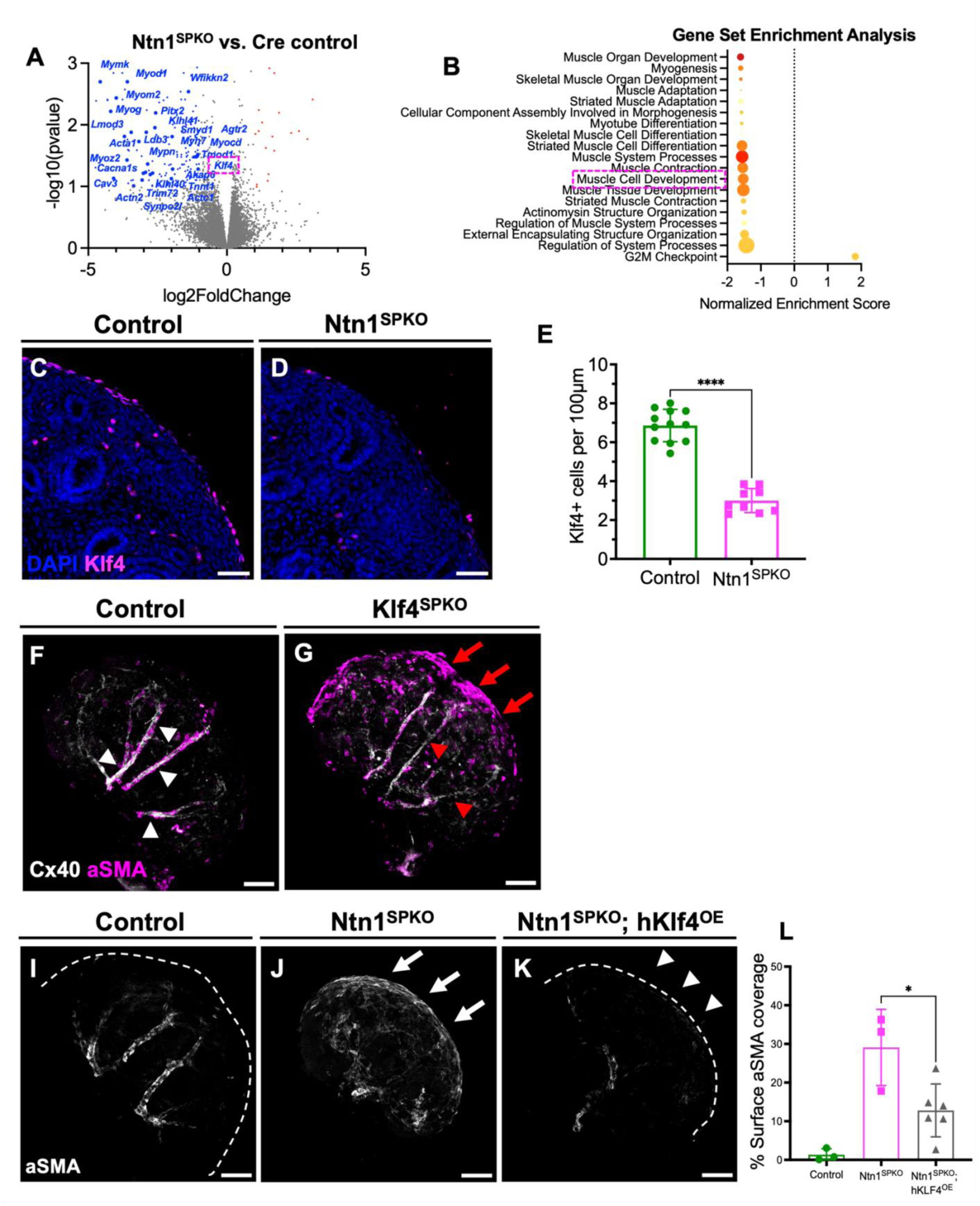
*Ntn1^SPKO^*kidneys display loss of Klf4 at E13.5 in stromal progenitors and premature differentiation into smooth muscle at the kidney cortex. (A) Volcano plot of differentially expressed genes. Red and blue dots indicate genes with at least 2-fold up- or downregulation and false discovery rate <0.1, respectively. Annotated genes are from GO:005501 (muscle cell development) or have known roles in muscle development. (B) Gene set enrichment analysis of differentially expressed genes, showing dysregulation of multiple processes involved in muscle cell development and differentiation. Size of dot correlates to the size of the gene set. (C,D) Immunofluorescence of Klf4 protein on E13.5 kidney sections, showing enrichment of Klf4 in cortical stromal progenitors and decrease in Klf4+ nuclei in *Ntn1^SPKO^* kidneys. (E) Quantification of nuclear Klf4+ foci along the surface of the kidney in control and *Ntn1^SPKO^* kidneys, showing a significant decrease (n=9, p<0.0001). (F-G) WMIF of E13.5 *Klf4^SPKO^* kidneys, showing decrease in arterial smooth muscle coverage (red arrowheads) and increase in smooth muscle cell differentiation at the kidney surface (red arrows). (I-K) WMIF for aSMA of E13.5 kidneys from control, *Ntn1^SPKO^*, or hKlf4 overexpression in *Ntn1^SPKO^* background, showing decreased surface smooth muscle upon overexpression of Klf4. Kidney surface is outlined where not visible. White arrows indicate significant smooth muscle coverage, white arrowheads indicate less smooth muscle upon overexpression of Klf4. (L) Quantification of percentage of surface covered by smooth muscle, measured by percent of pixel area in thresholded max intensity projections (n=3, p=0.0215). Arterial smooth muscle was discounted in quantifications by manually removing pixels from individual slices. Scale bars: 50µm (C-D), 100µm (F-K).

KLF4 is an evolutionarily conserved zinc-finger transcription factor that is required for maintenance of a mesenchymal state in vSMCs (Yap et al., 2021). KLF4 protein prevents binding of the muscle factor MYOCD, along with the transcriptional coactivators SRF and MRTF-A/B, to CarG boxes (Liu et al., 2005). In situ hybridization of developing kidneys showed expression of Klf4 in cortical stromal progenitors **(Fig. S6A’).** In situ data from earlier and later time points suggest that this expression is tightly temporally restricted and transient, as expression in the kidney periphery is low or not detectable at 11.5, nor at E15.5 **(Fig. S6A,A”,A’’’)**. Immunostaining for Klf4 protein at E13.5 revealed expression in Foxd1+ stromal progenitors (**Fig. 6C**).

We and others have identified previously uncharacterized molecular heterogeneity in the developing renal stromal progenitors using single cell RNA sequencing (scRNA-seq) (Combes et al. 2019, England et al. 2020). This is further evidenced here by expression of Klf4 in some, but not all, Foxd1+ stromal progenitors. RNAScope analysis of Ntn1, Foxd1, and Klf4 **(Fig S6B-B’’)** showed that most cells expressing Ntn1 were mutually exclusive from those expressing Klf4, suggesting that Klf4-expressing smooth muscle progenitors do not autoregulate via secretion of Ntn1 **(Fig. S6C)**. Indeed, Klf4 appears to mark the most peripheral cells in the developing E13.5 kidney, overlapping in expression with Foxd1, but not Ntn1 **(Fig. S6C)**. We note that we also see weak immunostaining in Sox17+ arterial endothelial cells **(Fig. S6D-D”)**. Klf4 is known to be involved in the response of arterial endothelial cells to blood flow, including initiating transcription of flow-responsive genes such as Cx40 (Chong et al., 2011). However, immunofluorescence of Klf4 in endothelial cells at this timepoint of development was low, suggesting a primary role in cortical stromal cells. Given the reported role of Klf4 in smooth muscle cells and the origin of smooth muscle cells within the Foxd1+ lineage, we infer that Klf4+ cells are precursors to renal vSMCs.

To confirm that loss of stromal Klf4 was occurring in *Ntn1^SPKO^* kidneys, we performed immunostaining for Klf4 in E13.5 control and *Ntn1^SPKO^* kidneys. In *Ntn1^SPKO^* kidneys, KLF4 was significantly reduced in the stromal progenitors (**Fig. 6C-E**). This decrease in Klf4 expression suggested a possible role in stromal and smooth muscle differentiation.

To assess whether Klf4 indeed plays a role in kidney development and test for a requirement for Klf4 in stromal progenitors, we ablated *Klf4* using the *Foxd1^GC^* driver line (*Klf4^SPKO^)*. Loss of *Klf4* from the stromal progenitors resulted in an accumulation of smooth muscle at the surface of the kidney, which was reminiscent of that seen in *Ntn1^SPKO^* **(Fig. 6F, G)**. While there remained scattered aSMA+ cells along distinguishable arteries in the *Klf4^SPKO^* mutants **(Fig. 6G**, red arrowheads), ectopic aSMA+ cells accumulated at the kidney periphery **(Fig. 6G**, red arrows). Notably, loss of stromal Klf4 did not result in defects in arterial patterning, suggesting that lack of mural cells was not solely responsible for the severe defects seen in *Ntn1^SPKO^* kidneys.

Finally, to show whether loss of Klf4 was directly responsible for ectopic smooth muscle differentiation at the surface of the kidney, we used a Cre-driven doxycycline-inducible overexpression system to selectively overexpress Klf4 in Foxd1-derived stromal cells in the *Ntn1^SPKO^* background. Upon expression of Klf4 in stromal progenitors, we observed significantly fewer vSMCs at the kidney surface **(Fig. 6I-L)**, suggesting that upon reintroduction of Klf4 expression, the ectopic smooth muscle caused by loss of Ntn1 had re-attained their mesenchymal fate and did not remain trapped at the kidney periphery.

## DISCUSSION

Stromal progenitors have been implicated in the patterning of the kidney and its vasculature, but how they mediate this crosstalk has been unclear. In this study, we identify a novel role for netrin-1 (Ntn1) in the early development and patterning of the kidney vasculature, as well as in proper localization of vascular mural cells. We show that profound defects in arterial patterning occur upon loss of Ntn1 expression from Foxd1+ stromal progenitor cells. These vessel defects are accompanied by impairment of mural cell recruitment to developing arteries. In addition, we show that stromal progenitors at the cortex of *Ntn1^SPKO^* kidneys prematurely and ectopically differentiate into contractile smooth muscle. We also find that the transcription factor Klf4 is significantly downregulated upon loss of Ntn1 in a non-cell autonomous manner, and that stromal loss of Klf4 leads to similar dysregulation of mural cell localization. Together, these data suggest that the Ntn1-Klf4 axis is a novel mediator of stromal crosstalk that is required for proper timing of mural cell differentiation and for proper vascular patterning within the developing kidney, underscoring the idea of an instructive and heterogeneous kidney stroma.

### Restriction of Ntn1 expression to the cortical stroma

Using single cell RNA sequencing and *in situ* hybridization, we and others identified Ntn1 expression in the cortical stromal progenitors during renal development (Combes et al., 2019; England et al., 2020). Our data shows that Ntn1 protein and RNA expression begins early in nephrogenesis. We show that Ntn1 is expressed by a subset of Foxd1+ stromal progenitors at E13.5, which is in line with our previous reports on the heterogeneity of the E18.5 stromal progenitors (England et al., 2020). By contrast, as a secreted ligand, Ntn1 protein is observed more diffusely within the cortex of the kidney and surrounding most of the stromal and epithelial cells.

Closer analysis at multiple timepoints shows that Ntn1 protein is always restricted to nephrogenic zone, and that protein appears to be enriched at the terminal tips of the ureteric bud. We posit that the UB tip may serve as a ligand sink that might further constrain Ntn1 protein presence to the most cortical regions. However, given the decrease in ureteric bud branching in *Ntn1^SPKO^* kidneys, a direct role cannot be excluded.

By P3, Ntn1 protein is no longer detectable in the kidney cortex, coinciding approximately with the cessation of nephrogenesis. This tight spatiotemporal restriction of Ntn1 RNA and protein is striking and suggests a specialized role in regulating kidney development. Given its restricted expression within a region relatively devoid of blood vessels, and its known role as a guidance cue for endothelial cells, we sought to test whether Ntn1 might play a role in determining where and when kidney blood vessels form.

### Ntn1 is essential to normal vascular patterning and arteriogenesis

Loss of Ntn1 from stromal progenitor cells results in profound defects of renal blood vessel development. One of the most striking defects in the *Ntn1^SPKO^* kidneys is abnormal arterial differentiation. Expression of the flow responsive gap junction protein Cx40 is decreased significantly within the kidney arteries. Normally, vessels connect to the kidney at two main entry points at the hilum, and branch thereafter at regular intervals, eventually connecting along arcuate arteries. This pattern is disrupted in the absence of Ntn1.

We find that the relatively stereotyped pattern of arterial branching is lost in Ntn1 mutants, which undoubtedly leads to defects in vascular perfusion. We propose that due to a combination of improper invasion of renal artery branches from the surface and abnormal collateral connections in the hilar and extrarenal regions, flow is shunted away from arteries within the kidney, resulting in failure of remodeling and consequent arteriogenesis. As hemodynamic flow and vessel remodeling is known to be essential for arteriovenous differentiation and for Cx40 expression, it is not surprising that abnormal vascular architecture in *Ntn1^SPKO^* kidneys may impair arteriogenesis. Indeed, we also observe decreased vessel perfusion in the absence of Ntn1 (data not shown). We speculate that defects in the architecture of the vasculature may shunt blood flow away from the main renal arteries, resulting in decreased flow and Cx40 expression. An alternative possibility is that the arterial identity in *Ntn1^SPKO^* kidneys is directly affected by the loss of Ntn1, however analysis of kidney specific Unc5b deletion will be needed to determine this.

In addition to defects in arteriogenesis and vessel disruption within the kidney, we also see gross vessel patterning defects at the hilum of the kidney and around the periphery of the kidney. Some of the defects observed, such as contribution of multiple aortic branches to the renal vasculature, are reminiscent of the vasculature observed in the global Foxd1 deletion mouse model (Hum et al., 2014, Sequeira-Lopez et al., 2015). These observations raise the possibility that ectopic extra-kidney vessels are not solely due to loss of cortical Ntn1, but to the compound loss of Ntn1 and the loss of one Foxd1 allele due to the *Foxd1^GC^.* The overlap of phenotypes could also suggest that Ntn1 is secreted around the periphery of the kidney, where it might be required for proper arterial patterning outside of the developing kidney.

While we find that overall corticomedullary patterning of arteries is restored as development continues, **Honeycutt et al**. in this issue perform a detailed analysis of arterial branching and area into adulthood and observe continued defects in *Ntn1^SPKO^* kidneys.

### Vascular smooth muscle cell (vSMC) loss and mislocalization in the absence of Ntn1

An important finding from our work and the work of others (**Honeycutt, et al**) is the decrease in smooth muscle coverage of developing renal arteries in *Ntn1^SPKO^* kidneys. One possibility is that vSMC loss is secondary to a defect in arteriogenesis and to a loss of endothelial secretion of chemotactic cues. However, genes such as *PDGFB* and *Pbx1*, both important cues for mural cell recruitment to arteries (Hurtado et al., 2015), were not found to be dysregulated by our RNA sequencing (see supplementary information). In addition, the altered differentiation and cell fate of smooth muscle progenitors and NG2+ cells at the kidney surface suggests an intrinsic defect within the stroma, rather than a secondary migration defect.

Recruitment of mural cells profoundly impacts vascular integrity and vascular function, and has more recently been implicated in vascular patterning (Armulik et al., 2011; Gaengel et al., 2009; Kemp et al., 2022; Orlich et al., 2022; Stratman et al., 2017). In this study, we show that ablation of a renal stromal factor, Ntn1, influences behavior and differentiation of the renal mural cells. We propose that blood vessels fail to properly pattern due to a non-autonomous effect. However, given that one of the Ntn1 receptors, Unc5b, is expressed in vascular endothelial cells, we cannot omit the possibility that Ntn1 from the stroma affects blood vessels directly.

### Stromal Ntn1 is required for normal kidney development

In addition to defects in arterial patterning and mural cell recruitment in *Ntn1^SPKO^* kidneys, we observe gross defects in the timing and progression of nephrogenesis. It is unclear whether these effects are due directly to a loss of Ntn1 signaling to epithelial cells or indirectly via other cell types such as the vasculature (either the endothelium or the mural cells). The extended presence of Six2+ nephron progenitor cells in Ntn1 mutants is mirrored by the disappearance of Six2+ cells, while Ntn1 expression is normally lost in normal kidneys. Hence, the question remains open as to how loss of Ntn1 impacts developing nephrons in the kidney.

Ntn1 could signal directly to epithelial cells. The Ntn1 receptor Neogenin is broadly expressed in the kidney, DCC is not detectable (data not shown) and Unc5c is expressed in nephron epithelium (Combes et al., 2019 and public datasets such as the Kidney Cell Explorer https://cello.shinyapps.io/kidneycellexplorer/ from Ransick et al., 2019). However, while global deletion of Unc5c impacts migration of axons, it does not appear to appreciably impact epithelial growth, as per **Honeycutt et al** in this issue. In addition, studies of Ntn1 have suggested it has signaling functions that are independent of known receptors (Wilson et al., 2006), underscoring the difficulty of identifying responsible receptors in the kidney.

It is worth noting that despite the prolonged expression of Six2, we observe fewer glomeruli in Ntn1 knockout kidneys, even when accounting for kidney weight/body weight ratio. This, along with analysis of other markers of nephron differentiation such as NCAM, suggests that prolonged nephrogenesis in the mutants is unproductive as *Ntn1^SPKO^*kidneys are markedly smaller. Another, perhaps more remote possibility, is that physical compaction of the kidney due to excess contractile smooth muscle cells at the developing kidney periphery results in physical suppression of branching of the ureteric tree. Further work will be required to identify possibly direct roles of Ntn1 in the developing kidney, however this will prove challenging given the complexity of the Ntn1 ligand relationship with its many different receptors, both known and unknown.

### Mural cell normalization at late gestation in the absence of Ntn1

Despite premature differentiation of smooth muscle progenitors and retention at surface of the kidney, we observe eventual restoration of mural cells around arteries in kidneys lacking Ntn1. This could indicate that some vSMC progenitors at the cortex in Ntn1 mutant embryos remain able to expand and differentiate into vascular smooth muscle, later associating with arteries. Indeed, we see some association of NG2+/aSMA-mural cells to arteries in *Ntn1^SPKO^* kidneys. In control kidneys, NG2+ cells surround inner layers of vSMCs, which co-express aSMA and NG2. Whether these cells are immature smooth muscle cells or a separate lineage capable of transdifferentiation remains to be seen. Alternatively, a separate source of smooth muscle progenitors could compensate for the inability of the Foxd1+/Klf4+ smooth muscle progenitors at the surface to invest along arteries. We favor this possibility, as we note that when looking at both aSMA and NG2+ mural cells, staining is stronger in vessels originating from outside the hilum and entering the kidney via the cortex compared to arteries fully within the kidney. This suggests that these ectopic vessels may be populated with mural cells originating from outside the kidney, such as aortic smooth muscle. Another potential source of alternative vSMC progenitors within the kidney is the Tbx18+ cell population arising from the ureteric stroma. These cells have been suggested to contribute to vSMCs in the kidney (Airik et al., 2006), but their contribution to the renal stroma is still poorly understood.

### Ntn1 is required in stromal progenitors for the muscle regulator Klf4

Vascular smooth muscle cells have been shown to originate from the cortical Foxd1+ stromal progenitors (Sequeira-Lopez et al., 2015). Here, we suggest they may arise from a subset of progenitors marked by Klf4 expression early in nephrogenesis. Klf4 has been shown to repress the activity of Myocardins along with their co-factors Srf to prevent or reverse smooth muscle differentiation, and it is often dysregulated during disease (Yap et al., 2021). We recently showed that stromal Srf was necessary for smooth muscle cell differentiation in the kidney (Drake et al, 2022). Given the known role of Klf4 in repressing Srf activity/smooth muscle cell differentiation and promoting a smooth muscle progenitor-like state (Majesky et al., 2017), we infer that Klf4 prevents mural cell progenitors from differentiating into smooth muscle within the cortical stroma. Loss of Klf4, due to loss of Ntn1, results in premature expression of smooth muscle genes in cortical progenitors and may block their ability to migrate and integrate into the periarterial niche, resulting in the ectopic localization of aSMA+ cells at the surface rather than around arteries. In addition, precocious differentiation of smooth muscle cells upon loss of Klf4 would explain the relative paucity of vSMC that we observe.

We note that our work does not distinguish whether loss of Ntn1 and Klf4 directly impacts either the ability of mural cells to migrate, or their ability to associate with endothelial cells within the perivascular niche (wrap around them). Given that we observe a decrease of mural cell investment along vessels, it is conceivable that either of these properties is defective. We posit that stromal cells at the kidney cortex are normally incorporated as the kidney grows and expands in size past them, meaning cortical regions eventually become more medullary regions. Hence, it is possible either migration or association could be hindered in in *Ntn1^SPKO^* kidneys. Future studies will be required to track migration of individual Klf4+ cells so see if that process is hindered in the absence of Ntn1, and to distinguish between these possibilities.

Furthermore, whether Ntn1 directly induces expression of Klf4 in smooth muscle progenitors or acts indirectly via another cell type to regulate smooth muscle differentiation remains unclear. We observe Neo1 expression within the cortical stroma by *in situ* hybridization, but given the heterogeneity of the stromal progenitors that we observe as well as the relatively ubiquitous expression of Neo1, further studies with higher resolution expression data are required to investigate the potential induction of Klf4 by Ntn1. Alternatively, netrin-1 could affect smooth muscle progenitors indirectly via a relay mechanism, either via other stromal cells or via endothelial cells (see **Fig. 7** model).

**Figure 7.**
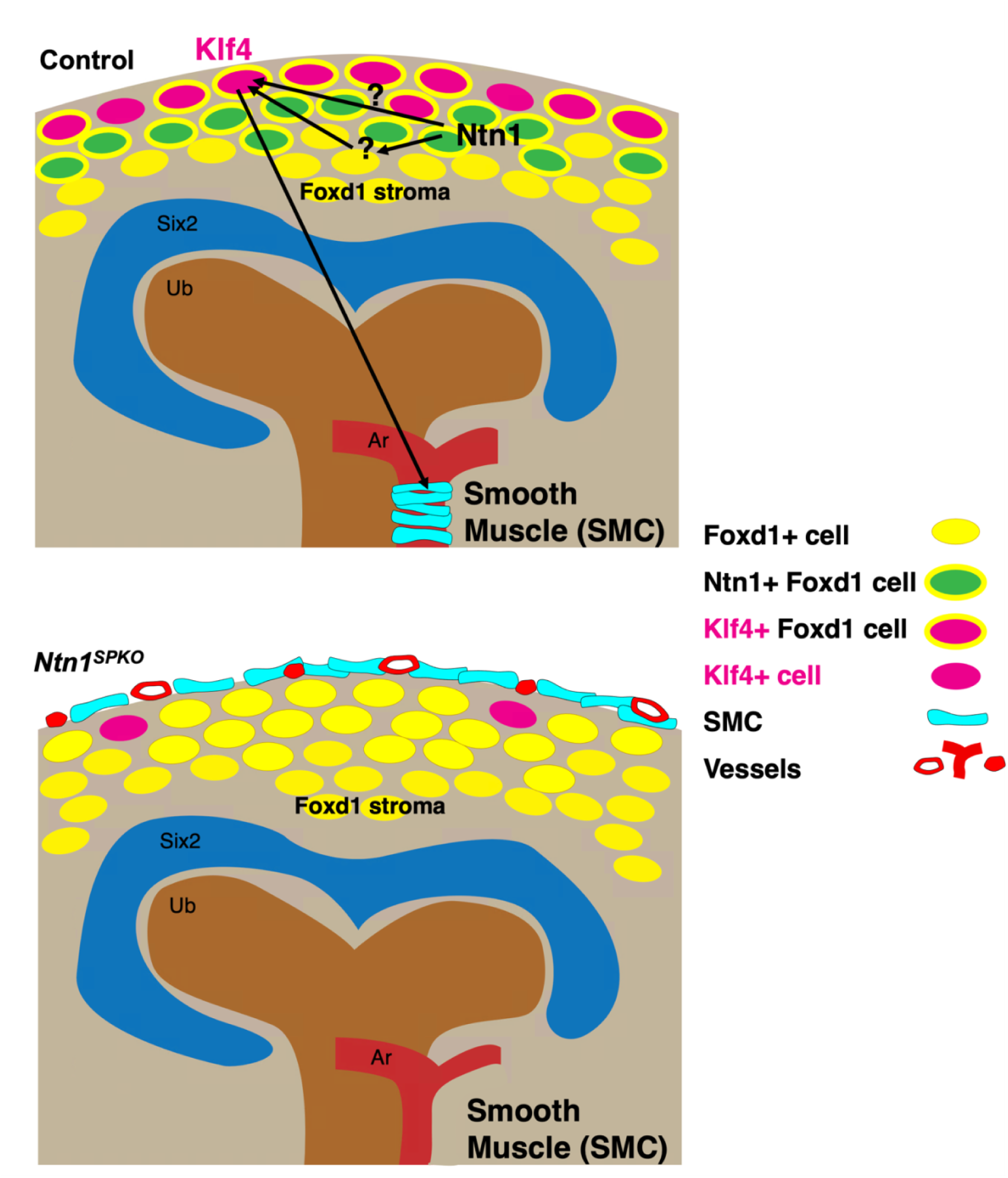
Model for mechanisms of netrin-1 mediated crosstalk. Ntn1 and Klf4 are expressed in different subpopulations of Foxd1+ cortical stromal progenitors. Without Ntn1 expression, there is decreased Klf4 expression within peripheral progenitors, which results in abnormal differentiation into contractile smooth muscle at the kidney periphery and decrease in arterial smooth muscle. Ntn1 may directly induce Klf4 expression or affect expression via an intermediate other cell type, such as other stromal progenitors, epithelial cells, or endothelial cells.

These results argue that Klf4 is required for stromal progenitors to remain in a mesenchymal and undifferentiated state and allow them to associate with the artery prior to their differentiation. Absence of Ntn1, and thus Klf4, leads to inappropriate and ectopic differentiation of smooth muscle progenitors, essentially decoupling smooth muscle differentiation and investment to arteries.

### Summary

In sum, these novel findings demonstrate how critical the kidney stroma is to overall kidney development. Restricted expression of a single guidance cue, Ntn1, is required for proper differentiation of three important lineages within the kidney: the stroma itself (its derivates, the mural cells), the vascular endothelium, and the nephron epithelium. Mechanistically, we show that a key muscle progenitor transcription factor Klf4 depends on Ntn1, and that loss of either of these factors impairs both mural cell investment and vascular development. Together, these findings further underscore that the stroma is not merely supportive of the organs it pervades, but rather it is a critical partner in successful organogenesis.

## MATERIALS AND METHODS

### Mouse Models

Experiments were performed in accordance with protocols approved by the University of Texas Southwestern Medical Center (UTSW) Institutional Animal Care and Use Committee. Timed matings were set up between females 7-9 weeks and males older than 7 weeks, and mating was ascertained by checking plugs, at which point the pregnancy was determined to be 0.5 days gestation. Dams were euthanized at desired timepoints and embryos were dissected for relevant tissues. The following mouse lines were used: *Foxd1^GC^* (JAX #:012463), Ntn1^flox^ (JAX#:028038), Klf4^flox^ (MMRRC 029877-MU), LSL-rTTA (Gift from Hao Zhu, UTSW. JAX # 029617), tet-on-hKLF4 (Gift from Mark Kahn, Penn. JAX# 019039). B6.Cg-Gt(ROSA)26Sortm14(CAG-tdTomato)Hze/J (JAX #:007914).

Klf4 overexpression was achieved in the *Ntn1^SPKO^* background using tet-on-hKLF4 and LSL-rTTA by providing doxycycline hyclate (Sigma, 150mg/mL) in drinking water along with 1% sucrose w/v from E11.5 until dissection (E13.5).

### Tissue preparation and immunofluorescence on sections

E13.5-P5 kidneys were dissected and fixed in 4% PFA/PBS overnight at 4 C, embedded in paraffin, and sectioned as previously described (Daniel et al., 2018). Briefly, tissue was dehydrated to 100% ethanol, incubated in xylene twice for 10 min, a mixture of 1:1 xylene and paraplast at 10 min at 60°C, and replaced with paraplast 3 times at 60C before embedding and sectioning at 10µm with a Leica microtome onto SuperfrostPlus glass slides.

Immunofluorescence staining was performed as previously described. Briefly, paraffin sections were deparaffinized with xylene, rehydrated through an ethanol series into PBS, permeabilized with 0.3% PBS-TritonX100 for 10 minutes with rotation, and treated with R-Buffer A or B for heat mediated antigen retrieval of nuclear and cytoplasmic antigens, respectively, in a 2100 Retriever (Electron Microscopy Sciences). Slides were allowed to cool and blocked in CAS Block (Invitrogen) for 2h at room temperature. Slides were incubated in primary antibody overnight at 4°C, washed in PBS and incubated in secondary antibody for 1.5 h at room temperature. Slides were washed with PBS, treated with a blood lysis buffer (10mM CuSO_4_, 50mM NH_4_Cl pH=5), incubated in deionized water for 5 min, and washed with PBS again before mounting with DAPI-Fluoromount (Southern Biotech). Imaging was performed on a Nikon A1R confocal microscope. Antibodies used, concentrations, and conditions are summarized in Table S1.

### Whole mount immunofluorescence (WMIF)

E13.5-E18.5 kidneys were dissected as above, ensuring that all surrounding mesenchyme was removed to limit extra-renal fluorescent signal with certain antibodies. Tissue was dehydrated into 100% MeOH and stored at 4C until staining. For WMIF, kidneys were rehydrated into PBS, permeabilized with 1% (E13.5) or 3% (E15.5, E18.5) TritonX100 in PBS for 3 hours (E13.5, E15.5) or 6 hours (E18.5). Kidneys were then blocked with CAS block for a minimum of 2 hr and incubated with primary antibody overnight at 4°C. Tissue was washed with PBS for a minimum of 6 hour-long washes, and incubated in secondary antibody overnight at 4°C. Tissue was washed again at least 6 times, dehydrated into 100% MeOH and cleared using 2:1 benzyl alcohol:benzyl benzoate (BABB) for at least 10 min. Cleared tissue was mounted in BABB into concavity slides (Electron Microscopy Sciences) and imaged using LSM700 Zeiss confocal microscope.

### Image analysis

Linear adjustments, maximum intensity projections, and cropping was performed in ImageJ. Manual removal of fluorescent signal was limited to ureteric smooth muscle and was performed slice by slice and indicated in figure captions. aSMA signal attributable to ureteric smooth muscle was easily distinguished from arterial smooth muscle. 3D projection and orthogonal projections were visualized using Imaris 9.0.0.2 or Imaris 10.0 software (Bitplane).

### In situ hybridization

Ntn1, Unc5b, Neo1, and Unc5c coding sequences were obtained from Dharmarcon (accession no: BC029161 – Ntn1, BC048162 – Unc5b, BC054540 – Neo1, BC115772 – Unc5c) and digoxigenin labeled RNA probes were synthesized as previously described (Daniel et al, 2018). Briefly, plasmid was linearized and purified by phenol:chloroform extraction. Probes were synthesized at 37°C for 2–4 hrs in digoxigenin-synthesis reaction mixture with T7 RNA polymerase (Roche). After synthesis, DNA was eliminated with DNase I (Promega) and RNA probes were purified using Micro Bio-Spin columns (Bio-RAD). Final concentration for hybridization was 1 µg/mL and was achieved by diluting in pre-hybridization buffer. *In situ* hybridization was performed on paraffin embedded sections as previously described (Daniel et al, 2018). Briefly, slides were deparaffinized in xylene and rehydrated, after which they were treated with 15 µg/mL Proteinase K for 15 min and fixed in 4% PFA. Slides were incubated in prewarmed pre-hybridization buffer for 1 hour followed by overnight incubation with diluted RNA probe at 65°C. Slides were then washed in 0.2x SSC and MBST before blocking with 2% blocking solution (Roche) for at least 1 hour at room temperature. Slides were then incubated overnight with Anti-Dig alkaline phosphatase-conjugated antibody (Roche, 1:4000) at 4°C. Slides were washed with MBST and NTMT and incubated with BM Purple (Roche) for colorimetric reaction. Slides were post-fixed in 4% PFA and mounted in Permount (Fisher). Slides were imaged with Zeiss Axiovert 200M scope.

### Western blot

E13.5 kidneys were dissected as above. Protein was extracted in RIPA buffer for 30 minutes and quantified by BCA assay (Thermo). 10 µg protein was run on a polyacrylamide gel, blocked with blocking buffer (Bio-RAD), blotted with primary antibody overnight at 4°C, washed with TBST, incubated in secondary antibody, and developed using ECL and x-ray film. Densitometry analysis was performed on scanned images using Fiji.

### RNAscope fluorescent *in situ* hybridization

RNAscope probes were obtained from ACD (Mm-Klf4 (426711), Mm-Ntn1-C2 (407621-C2), Mm-Foxd1-C3 (495501-C3)). E13.5 kidneys were dissected as above and fresh frozen in OCT compound (Scigen) with liquid nitrogen. Kidneys were sectioned with a cryostat at 10µm and RNAscope fluorescent multiplex hybridization was performed following the manual using the multiplex reagent kit (320850). Images were acquired on a Nikon A1R confocal microscope. ROIs were created in ImageJ based on DAPI counterstain using the Analyze Particles tool, and RNAscope dots within each nucleus were counted manually. Cells were classified based on how many RNAscope dots of each gene were present. Only cells with Foxd1 expression were included in analysis. Cells with only one of Ntn1 or Klf4 dots, in addition to cells with a much higher proportion dots from one gene (over 2 times as many), were counted as being Foxd1/Ntn1+ and Foxd1/Klf4+, respectively. Cells with similar number of Ntn1 and Klf4 dots were counted as triple positive Foxd1/Ntn1/Klf4+ (less than 2 fold different).

### Glomerular count (Acid maceration)

HCl maceration of whole kidneys were performed according to MacKay et al., (1987). Briefly, kidneys were isolated from 6 week-old control and *Ntn1^SPKO^* mice, decapsulated, roughly chopped with a razor blade and incubated in 5ml 6M HCl per kidney for 90min. Every 30min, the kidneys were pipetted up and down to further disrupt the kidneys. Digested kidneys were diluted with 5 volumes (25ml) of distilled water and incubated at 4°C overnight. For counting, 1 ml of macerate was pipetted into a cell culture dish with grid lines and glomeruli were counted.

### RNA sequencing and differential gene analysis

Transcript abundance was estimated without aligning reads using Salmon (Patro et al., 2017) against an index of coding sequences from the Ensembl GRCm38 assembly. Transcript-level abundance was imported and count and offset matrices generated using the tximport R/Bioconductor package. Differential expression analysis was performed using the DESeq2 R/Bioconductor package (Love et al., 2014). Gene sets were downloaded from MSigDB (Liberzon et al.; Liberzon et al., 2011; Subramanian et al., 2005) using the msigdbr R package. The R/Bioconductor package fgsea was used to carry out gene set enrichment analysis.

### Data analysis and visualization

All data was plotted and graphs were generated in PRISM 9 XML. All comparisons were done using the parametric unpaired Student’s t-test. ns=p>0.05; *=0.01<p<0.05; **=0.001<p<0.01; ***=0.0001<p<0.001; ****=p<0.0001. Linear manipulations of images were done using ImageJ. Figures and models were made using Microsoft Powerpoint, BioRender.com, and Adobe Illustrator.

## ACKNOWLEDGEMENTS

We thank the entire Cleaver lab for critical discussions and input on the manuscript.

We thank Mark Kahn for the tet-on-hKLF4 mouse line and Hao Zhu for the lsl-rTTA mouse line. We thank Lu Sun and his lab for help with the RNAscope protocol and analysis. We are grateful to Genepaint (gp3.mpg.de) and the Allen Brain Atlas for *in situ* hybridization data. We are also grateful to ongoing discussions with the O’Brien lab, as we developed our stories independently, but with helpful occasional crosstalk.

## FOOTNOTES

### Author contributions

Conceptualization: P.L., X.G., O.C.; Methodology: P.L., X.G., O.C.; Formal analysis: P.L., C.C., X.G.; Investigation: P.L., X.G., O.C; Resources: O.C..; Data curation: P.L., C.C., X.G.; Writing - original draft: P.L., O.C.; Writing - review & editing: P.L., X.G., T.J.C., O.C.; Visualization: P.L., X.G.; Supervision: O.C.; Project administration: P.L.; Funding acquisition: O.C.

